# A phytocannabinoid-sensitive phosphorylation switch converts endocannabinoids into alternative lipid GPCR activators

**DOI:** 10.64898/2026.09.21.753239

**Authors:** Anna C. Love, Yasuyuki Kihara, Hirotaka Mizuno, Kazufumi Nagai, David Coronel, Douglas J. Sheffler, Valerie P. Tan, Bradley S. Moore, Jerold Chun

## Abstract

Endocannabinoids (eCBs) like anandamide and 2-arachidonoylglycerol are endogenous lipid ligands for CB_1_ and CB_2_ G protein-coupled receptors (GPCRs). Here, we show that phosphorylation of biologically important anandamide generates naturally occurring anandamide phosphate (AEAp), which switches GPCR ligand specificity to lysophosphatidic acid (LPA) receptors (LPARs) and the primate-specific bile acid sensory receptor MRGPRX4. The kinase responsible for anandamide phosphorylation was identified as diacylglycerol kinase (DGK) theta (DGKθ or DGKQ) using activity-guided brain fractionation, inhibitor profiling, recombinant reconstitution, and DGK isozyme screening. DGKQ showed noncanonical biphasic lipid kinetics and was inhibited by phytocannabinoids, most notably tetrahydrocannabinolic acid. Thus, eCBs are not only cannabinoid receptor ligands but also enzymatically adaptable lipid signals activating LPARs and MRGPRX4, thus linking metabolism of the distinct lipid LPA with *Cannabis* pharmacology.

## Main Text

Bioactive lipids are produced transiently and activate diverse cellular responses through specific G protein-coupled receptors (**GPCRs**) (*1*). Among the earliest lipid GPCRs to be molecularly characterized was the cannabinoid receptor CB_1_, which was initially identified as a receptor for plant-derived cannabinoids (*e.g.,* phytocannabinoids) (*2*). Characterization of CB_1_ subsequently led to the discovery of its endogenous ligands, endocannabinoids (**eCBs**), including anandamide (**AEA**) and 2-arachidonoylglycerol (**2-AG**) (*3-5*). These eCBs regulate synaptic transmission, neuroinflammation, pain, and mood (*6, 7*). Unlike classical neurotransmitters, eCBs are synthesized on demand, requiring rapid metabolic turnover to achieve spatial and temporal control over signaling. Their transient signaling therefore depends on rapid enzymatic degradation, with AEA and 2-AG primarily hydrolyzed by fatty acid amide hydrolase and monoacylglycerol lipase, respectively (*8, 9*). Phytocannabinoids such as Δ ^9^-tetrahydrocannabinol (**Δ^9^-THC**) and cannabidiol (**CBD**) also modulate many of these physiological processes through mechanisms that extend beyond direct cannabinoid receptor activation (*10*). CBD has been reported to inhibit AEA hydrolysis and cellular transport, suggesting that phytocannabinoids can influence eCB signaling at the level of both receptor activity and metabolism (*10*).

Chemical modification can further alter eCB signaling. For example, oxidative metabolism generates bioactive eCB derivatives, including prostamides and prostaglandin glycerol esters (*11*). These metabolites exhibit biological activities distinct from their parent eCBs, highlighting how chemical modification expands the signaling repertoire of eCBs. Beyond these established transformations, phosphorylation is an emerging yet underexplored chemical modification of eCBs. The phosphorylated derivative of 2-AG is 2-arachidonoyl lysophosphatidic acid (2-arachidonoyl-**LPA**), a naturally occurring LPA species. LPA signals through six LPA receptors (**LPARs**; LPA_1-6_), with LPA_1-3_ clustering near cannabinoid receptors CB_1_ and CB_2_ in a phylogenetic analysis of rhodopsin-family GPCRs (**Fig. 1A**) (*12*). Similarly, phosphorylation of AEA would generate anandamide phosphate (**AEAp**), which was previously identified as a transient intermediate in an alternative biosynthetic pathway for AEA (*13*). Additionally, structural, binding, and pharmacological studies have implicated it as an LPA_1_ ligand (*14, 15*). Altogether, these observations suggest that AEAp may function as a GPCR-activating lipid mediator and raise the possibility that phosphorylation could serve as a molecular switch that redirects eCBs from cannabinoid to non-cannabinoid GPCR signaling. However, which enzyme is responsible for eCB phosphorylation, and whether phytocannabinoids modulate this enzyme, remains unknown. Here, we report the molecular identity of a phytocannabinoid-sensitive endocannabinoid kinase that directly phosphorylates eCBs to generate non-cannabinoid GPCR-activating phospho-eCBs, thereby extending eCBs into new central nervous system biology.

**Fig. 1.**
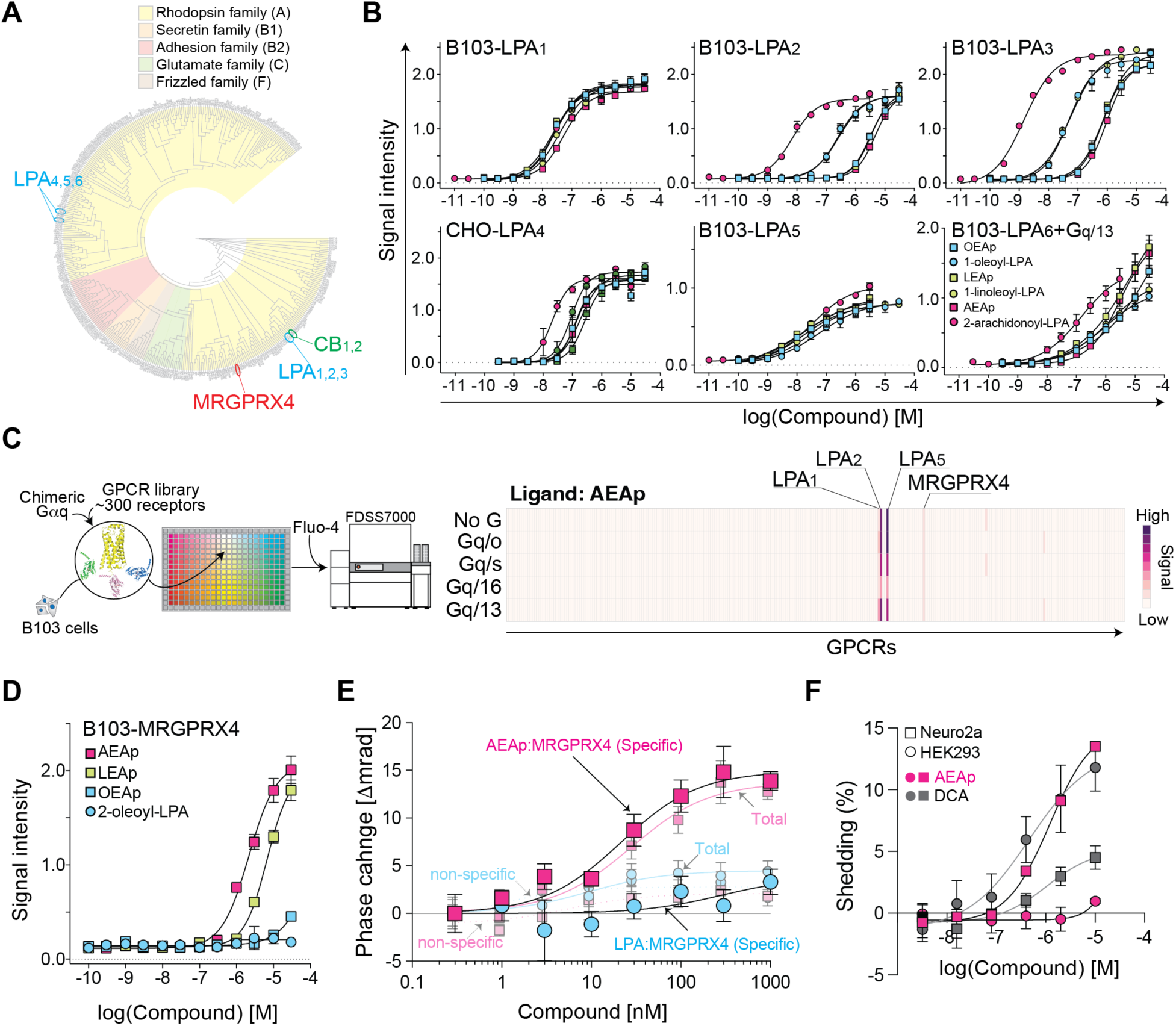
Phosphorylated endocannabinoid-related lipids define a GPCR-active mediator class. **(A)** Phylogenetic tree of human GPCRs showing the positions of LPARs, cannabinoid receptors, and MRGPRX4 within the rhodopsin-family GPCR landscape. **(B)** Intracellular Ca^2+^ mobilization in B103 or CHO cells heterologously expressing human LPARs treated with *N*-oleoylethanolamine phosphate (OEAp), *N*-linoleoylethanolamine phosphate (LEAp), anandamide phosphate (AEAp), 1-oleoyl-LPA, 1-linoleoyl-LPA, or 2-arachidonoyl-LPA. LPA_6_ responses were measured in B103-LPA_6_ cells co-expressing a Gα_13_-Gα_q_ chimeric G protein. EC_50_ values are provided in Table S1. **(C)** Ca^2+^ signaling-based rhodopsin-family GPCR screen for AEAp responsiveness in the absence or presence of chimeric G proteins. Robust responses were detected for LPA_1_, LPA_2_, and LPA_5_, and an additional response was identified at MRGPRX4. **(D)** Concentration-dependent Ca^2+^ mobilization in MRGPRX4-expressing B103 cells treated with AEAp, LEAp, OEAp, or 2-oleoyl-LPA. AEAp and LEAp activated MRGPRX4, whereas 2-oleoyl-LPA did not. **(E)** Backscattering interferometry analysis of AEAp and LPA binding to MRGPRX4. Specific binding was calculated by subtracting nonspecific binding from total binding. AEAp showed specific binding to MRGPRX4, whereas LPA showed no detectable specific binding. Data are mean ± SEM; n = 5. **(F)** TGF-α shedding assay in HEK293 or Neuro2a cells expressing MRGPRX4. Cells were treated with AEAp or deoxycholic acid (DCA) to compare cell-context-dependent MRGPRX4 activation. DCA activated MRGPRX4 in both cell types, whereas AEAp showed measurable activation only in Neuro2a cells. The EC_50_ values for DCA were 0.46 µM in HEK293 cells and 1.06 µM in Neuro2a cells; the EC_50_ value for AEAp in Neuro2a cells was 1.2 µM. Data are mean ± SEM.

## Results

### Phosphorylation converts eCB-related lipids into GPCR ligands

To test whether AEAp acts as a broad LPAR ligand, beyond the previously observed activity at LPA_1_ (*14, 15*), we performed intracellular Ca^2+^ mobilization assays in cells heterologously expressing a series of human LPARs (**Fig. 1B and Fig. S1A-C**). Cells were stimulated with AEAp, and related phosphorylated *N*-acylethanolamides (including *N*-oleoylethanolamine phosphate and *N*-linoleoylethanolamine phosphate), or corresponding LPA species with matched fatty acyl chains. All tested phosphorylated *N*-acylethanolamides activated LPARs with receptor-subtype selectivity and potency. The activity was particularly potent at LPA_1_ and LPA_5_, where responses approached those of canonical LPA ligands (**Fig. 1B and Table S1**). These results demonstrate that phosphorylation can convert AEA and other *N*-acylethanolamides into LPAR agonists.

AEA has extensive biological activity as documented in the nervous system (*6, 7*), suggesting that AEAp may serve as a ligand for additional GPCRs. Our screen identified the primate-specific sensory receptor MRGPRX4 as a new AEAp-responsive receptor (**Fig. 1C**). AEAp (and *N*-linoleoylethanolamine phosphate) activated MRGPRX4-expressing cells with estimated EC_50_ values of 2.2 and 6.7 µM, respectively (**Fig. 1D**), comparable to reported EC_50_ values (2.6 µM) for the bile acid ligand, deoxycholic acid (*16, 17*). Direct binding of AEAp to MRGPRX4 was further confirmed using backscattering interferometry, a label-free optical method that detects molecular interactions through refractive index changes, resulting in a K_D_ value of 21.5 nM (**Fig. 1E**) (*15, 18*). Structural modeling placed the phosphate group of AEAp near a positively charged arginine cluster within the receptor pocket (**Fig. S1D**) (*19*). Notably, the predicted phosphate-binding geometry closely resembles that observed in the MRGPRX4 structure bound to a synthetic, phosphorylated deoxycholic acid derivative (*20*). In contrast, LPA neither transduces Ca^2+^ signals nor showed specific binding (**Fig. 1D and E**), indicating that MRGPRX4 distinguishes AEAp from LPA.

Importantly, the apparent ligand selectivity depended on cellular context (**Fig. 1F**). To further assess MRGPRX4 activation, we used the TGF-α shedding assay, a cell-based assay measuring GPCR activation (*21*). This assay previously identified deoxycholic acid as an MRGPRX4 agonist (*16*), and we reproduced deoxycholic acid-dependent MRGPRX4 activation in HEK293 cells, whereas AEAp had little or no response (**Fig. 1F**). However, in Neuro2a cells, AEAp acted as a full agonist, while deoxycholic acid behaved as a partial agonist, producing a maximal response approximately one-third that of AEAp (E_max_ = 5.19 vs. 15.18). These results suggest that neuronal cellular context favors AEAp-dependent MRGPRX4 activation with relevance to nervous system activity. These findings indicate that phosphorylation of anandamide alters its receptor selectivity across distinct GPCRs.

### Activity-guided fractionation identifies DGKQ as an AEA kinase

AEAp has been described as an intermediate in an alternative biosynthetic pathway of AEA, in which phospholipase C releases AEAp from *N*-acyl-phosphatidylethanolamine, and subsequent phosphatase activity converts AEAp to AEA (**Fig. 2A**) (*13*). However, our receptor studies demonstrate that AEAp is a receptor-activating lipid mediator in addition to a biosynthetic intermediate. This raised the question of whether AEA itself could be *actively* phosphorylated to AEAp, creating a direct route to phospo-eCBs. This hypothesis is supported by prior reports that acylglycerol kinase can phosphorylate monoacylglycerols, including 2-AG, to generate LPA species (*22*).

**Fig. 2.**
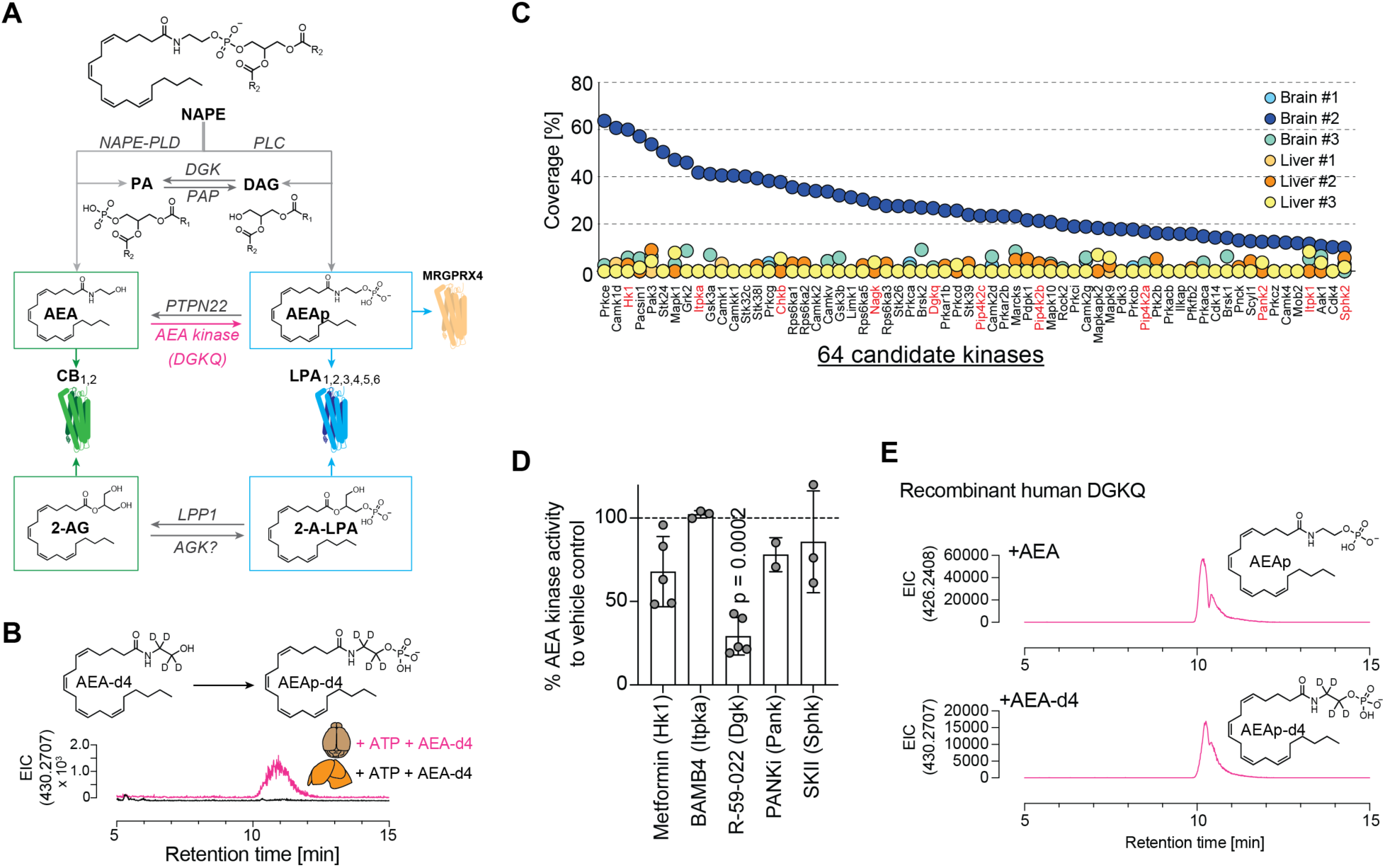
Activity-guided identification of DGKQ as an anandamide kinase. **(A)** Metabolic model linking endocannabinoid and LPA signaling pathways. 2-AG phosphorylation to 2-arachidonoyl-LPA (2-A-LPA) is shown as a parallel branch. The AEA-to-AEAp step is labeled as AEA kinase (DGKQ). **(B)** Detection of AEA kinase activity in mouse brain homogenates. Mouse brain and liver homogenates were separately incubated with deuterium-labeled AEA (AEA-d4) and ATP in the presence of phosphatase inhibitors, and the products were analyzed by LC-MS. AEAp-d4 was detected in brain homogenates in an ATP-dependent manner but was not detected in liver homogenates. **(C)** Candidate kinase prioritization by activity-guided fractionation and proteomics. Size-exclusion chromatography separated brain and liver homogenates into three fractions, and AEA kinase activity was detected only in brain fraction 2. Proteomic analysis identified 64 candidate kinases enriched in the active brain fraction relative to the inactive brain and liver fractions; kinases with known small-molecule or lipid-metabolite substrates are highlighted in red. **(D)** Pharmacological screening of candidate kinase classes in brain homogenate AEA kinase assays. The DGK inhibitor R-59-022 significantly reduced AEAp production, whereas inhibitors targeting other candidate kinase classes did not. p value was determined by two-way ANOVA. Data are mean ± SEM. **(E)** Recombinant human DGKQ directly phosphorylates AEA. LC-MS extracted-ion chromatograms (EIC) show production of AEAp from AEA and AEAp-d4 from AEA-d4 by recombinant human DGKQ in the presence of ATP and Mg^2+^, confirming that DGKQ can directly generate AEAp.

To test this possibility, mouse brain and liver homogenates were incubated with deuterium-labeled AEA and ATP in the presence of sodium orthovanadate to suppress phosphatase-mediated dephosphorylation. High resolution liquid chromatography-mass spectrometry analysis showed deuterium-labeled AEAp in brain homogenates, but not in liver homogenates (**Fig. 2B**). This activity was abolished by heat inactivation (**Fig. S2A**), indicating that AEAp production was enzymatic. The putative enzyme preferentially used ATP rather than GTP as the phosphate donor and required divalent cations, with Mg^2+^ supporting activity more effectively than Zn^2+^ (**Fig. S2B, C**). Thus, the mouse brain contains a unique ATP-dependent AEA kinase activity capable of generating AEAp directly from AEA.

Because 2-AG reduced AEAp production in brain homogenates (**Fig. S2D**), we considered whether the known monoacylglycerol kinase might be responsible for this activity. However, AEAp production was not enriched in crude mitochondrial fractions where acylglycerol kinase activity is localized (**Fig. S2E**). Furthermore, acylglycerol kinase cellular lysates did not phosphorylate either 2-AG or AEA under our assay conditions (**Fig. S2F**) (*22*). These data suggested that the observed brain AEA kinase activity was distinct from acylglycerol kinase (*22*). We therefore set out to identify the enzyme capable of generating AEAp.

Activity-guided fractionation of brain vs. liver homogenates followed by proteomic analysis was performed (**Fig. S2G**). The mass spectrometry assay showed AEA kinase activity only in one of three brain fractions, with no detectable activity in any liver fraction. The active brain fraction contained 871 proteins from which 11 kinase candidates with known small molecule substrate specificity were prioritized (**Fig. 2C**). Pharmacological screening of these candidates identified significant inhibition only with the diacylglycerol kinase (**DGK**) inhibitor R-59-022 (**Fig. 2D**), implicating DGK theta (DGKθ), also known as DGKQ (*23*), as the AEA kinase. DGKQ was originally cloned and characterized as the only type V DGK that phosphorylates diacylglycerol to produce phosphatidic acid (*24*).

Recombinant DGKQ phosphorylated AEA to AEAp (**Fig. 2E**). The biochemical preferences of recombinant human DGKQ recapitulated those observed in mouse brain homogenates, with DGKQ using ATP rather than GTP as the phosphate donor, requiring Mg^2+^ for activity, and showing maximal activity around pH 8.5 (**Fig. S3**). These results show that DGKQ/DGKθ (*23*) is an AEA kinase capable of actively generating AEAp directly from AEA.

### DGKQ functions as an eCB kinase

The human/mouse DGK family comprises ten lipid kinases that canonically phosphorylate diacylglycerol to phosphatidic acid, but differ substantially in domain architecture, regulation, subcellular localization, and substrate preference (*25*). To investigate whether AEAp production is specific to DGKQ or more promiscuous for the DGK family, each human DGK was overexpressed in HEK293 cells and assayed by mass spectrometry using multiple reaction monitoring. DGKQ was the only family member to produce AEAp significantly (**Fig. 3A**). Because 2-AG competed with AEA phosphorylation in brain homogenates (**Fig. S2D**), and monoacylglycerols have been reported as DGK substrates (*26*), we also tested 2-AG phosphorylation across the DGK family. In contrast to AEA, 2-AG phosphorylation was observed with several DGKs, with the strongest activities detected for DGKQ, followed by DGKA and DGKB (**Fig. 3B**). Thus, AEA is specifically phosphorylated by DGKQ, contrasting with 2-AG phosphorylation by multiple DGK enzymes.

**Fig. 3.**
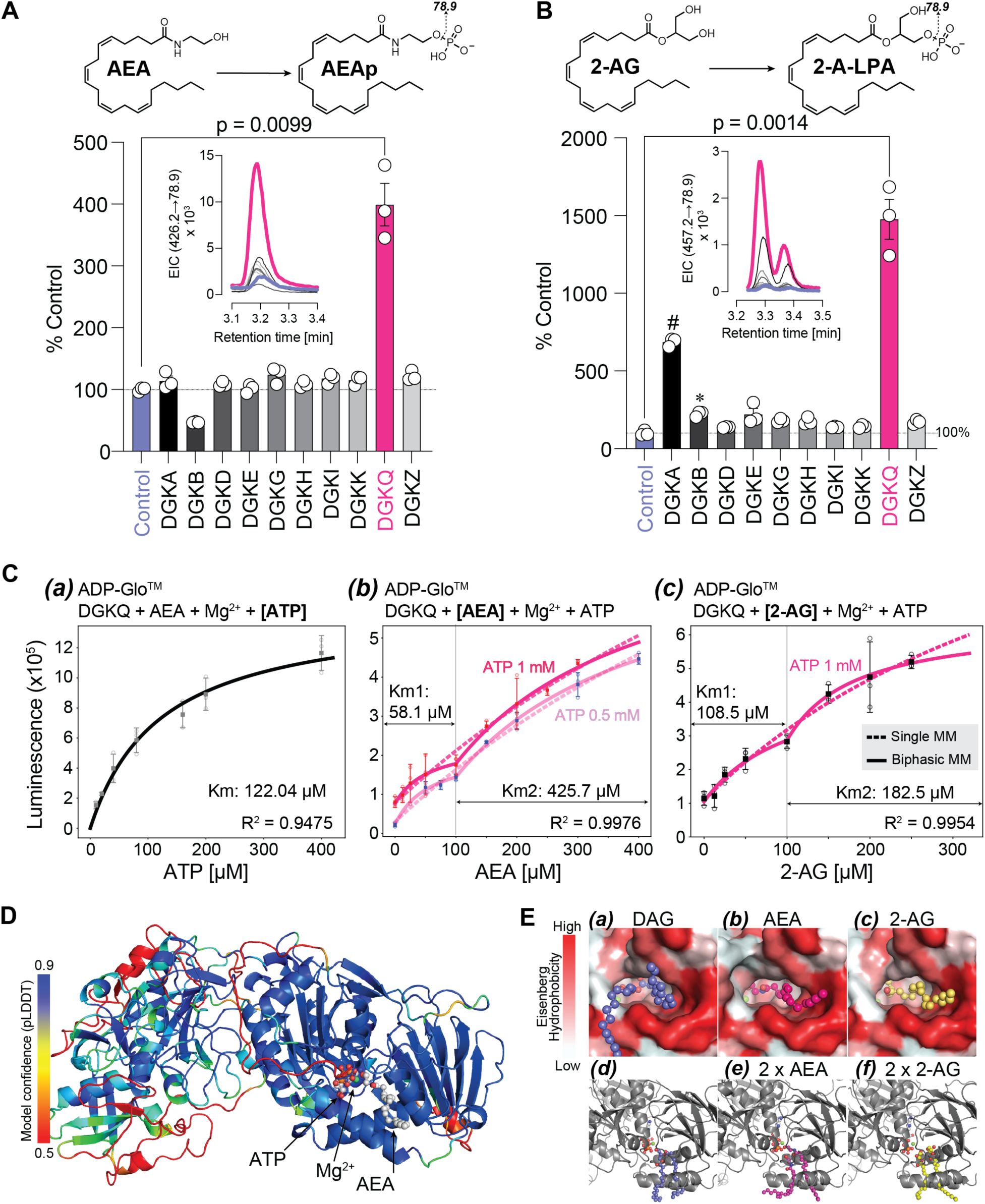
DGKQ is an endocannabinoid kinase that phosphorylates AEA and 2-AG. **(A)** AEA phosphorylation activity across the human DGK family. HEK293 cells expressing each DGK isozyme were incubated with AEA, and AEAp production was measured by LC-MS. DGKQ was the only DGK family member that significantly increased AEAp production. The inset shows multiple reaction monitoring chromatograms for AEAp detection using the 426.2→78.9 transition. P value was determined by two-way ANOVA. Data are mean ± SEM. **(B)** 2-AG phosphorylation activity across the human DGK family. HEK293 cells expressing each DGK isozyme were incubated with 2-AG, and 2-arachidonoyl-LPA production was measured by LC-MS. In contrast to AEA, 2-AG was phosphorylated by multiple DGK family members, with the strongest activity detected for DGKQ. The inset shows multiple reaction monitoring chromatograms for 2-arachidonoyl-LPA (2-A-LPA) detection using the 457.2→78.9 transition. p value was determined by two-way ANOVA. Data are mean ± SEM. **(C)** Enzymatic characterization of recombinant human DGKQ using the ADP-Glo^TM^ assay under non-micelle conditions. ***(a)*** ATP dependence of DGKQ activity with AEA as substrate, yielding an apparent ATP Km of 122.0 µM. ***(b)*** Concentration dependence of AEA on DGKQ activity at 0.5 or 1 mM ATP. Phosphorylation of AEA showed biphasic kinetics, with apparent Km values of 58.1 µM and 425.7 µM for the low- and high-concentration phases, respectively. ***(c)*** Concentration dependence of 2-AG on DGKQ activity. Phosphorylation of 2-AG also showed biphasic kinetics, with apparent Km values of 108.5 µM and 182.5 µM for the two phases. Solid lines indicate single Michaelis-Menten (MM) fits; dashed lines indicate biphasic MM fits. Model comparison by Akaike information criterion (AIC) favored the biphasic MM model (313.5) over the single MM model (333.6) for AEA phosphorylation (ΔAIC = 20.1). **(D)** Boltz-2 model of DGKQ bound to AEA, ATP, and Mg^2+^. The model is colored by predicted local distance difference test (pLDDT) confidence score. The catalytic region shows higher model confidence than the N-terminal regions and positions AEA near ATP and Mg^2+^ within the substrate-binding pocket. **(E)** Structural models of the DGKQ substrate pocket bound to DAG ***(a)***, AEA ***(b)***, 2-AG ***(c)***, or two monoacyl eCB substrates ***(d-f)***. DAG fills the hydrophobic pocket with two acyl chains ***(a, d)***, whereas AEA or 2-AG occupies one side of the pocket and leaves additional space ***(b, c)***. Co-folding with two AEA or two 2-AG molecules fills the pocket more completely ***(e, f)***, providing a potential structural rationale for a multiple-occupancy model of eCB phosphorylation.

DGK activity has classically been measured using a detergent-containing, lipid-micelle-based assay system (*26*). Because DGKQ-dependent AEA phosphorylation was detected without intentional micelle formation, we established a non-micellar ADP-Glo^TM^ assay to quantify DGKQ’s eCB kinase activity. Under these conditions, DGKQ showed an apparent ATP Km of 122 µM (**Fig. 3C**), which is consistent with previously reported ATP Km values for DGKs, typically in the 100∼500 µM range (*27, 28*). Interestingly, AEA and 2-AG phosphorylation did not conform to a single Michaelis-Menten relationship. Instead, both substrates showed biphasic kinetics, with a transition near 100 µM of substrate. For AEA, global fitting with a biphasic Michaelis-Menten model yielded apparent Km values of 58.1 µM and 425.7 µM for the two phases (**Fig. 3C**). This profile persisted independent of ATP concentration, indicating that the biphasic model was not due to ATP depletion, but rather reflected DGKQ’s specific kinetic properties. A similar biphasic profile was observed for 2-AG, with apparent Km values of 108.5 µM and 182.5 µM (**Fig. 3C**). High micromolar substrate Km values align well with other characterized lipid kinases. For example, ceramide kinase exhibits an apparent ceramide Km of ∼187 µM (*29*), while brain DGKs exhibit a diacylglycerol Km of ∼60 µM (*30*). Such biphasic features are consistent with non-Michaelis-Menten mechanisms previously observed in lipid processing enzymes and those with large hydrophobic active sites (*31*), where substrate partitioning, multiple substrate occupancies, or substrate-induced conformational states can alter apparent kinetics.

Structural modeling provided a potential explanation for the difference in observed features from DGKQ with monoacyl eCBs vs. diacylglycerol. Importantly, models of DGKQ co-folded with ATP, Mg^2+^, and either diacylglycerol, AEA, or 2-AG recapitulated the conserved ATP interaction with the glycine-rich GxGxxG motif, including G648 (*32*). This residue is required for DGKQ activity (**Fig. 3D**), validating the modeling outputs. In the model, diacylglycerol filled the hydrophobic pocket with two acyl chains and positioned its glycerol backbone toward the ATP-binding site (**Fig. 3E**). By contrast, AEA and 2-AG occupied the more hydrophobic side of the pocket, leaving space that could accommodate a second monoacyl substrate molecule (**Fig. 3E**). Co-folding with two AEA or 2-AG molecules filled this pocket more completely without disrupting ATP/Mg^2+^ positioning (**Fig. 3E and Fig. S4**). Together, these data identify DGKQ as an eCB kinase that converts AEA and 2-AG into the GPCR-active phosphorylated lipids AEAp and 2-arachidonoyl-LPA, respectively.

### Phytocannabinoids inhibit DGKQ to modulate the AEA-to-AEAp signaling switch

Given that phytocannabinoids influence eCB metabolism and receptor signaling (*33*), we tested whether they can also modulate the phosphorylation activity of DGKQ on eCBs (**Fig. 4A**). Both THC-rich and CBD-rich *Cannabis* extracts inhibited DGKQ-mediated phosphorylation of AEA in a concentration-dependent manner (**Fig. 4B**). These results suggest that DGKQ is sensitive to phytocannabinoids, expanding their bioactivity beyond direct cannabinoid receptor modulation to the regulation of eCB metabolism (*10*).

**Fig. 4.**
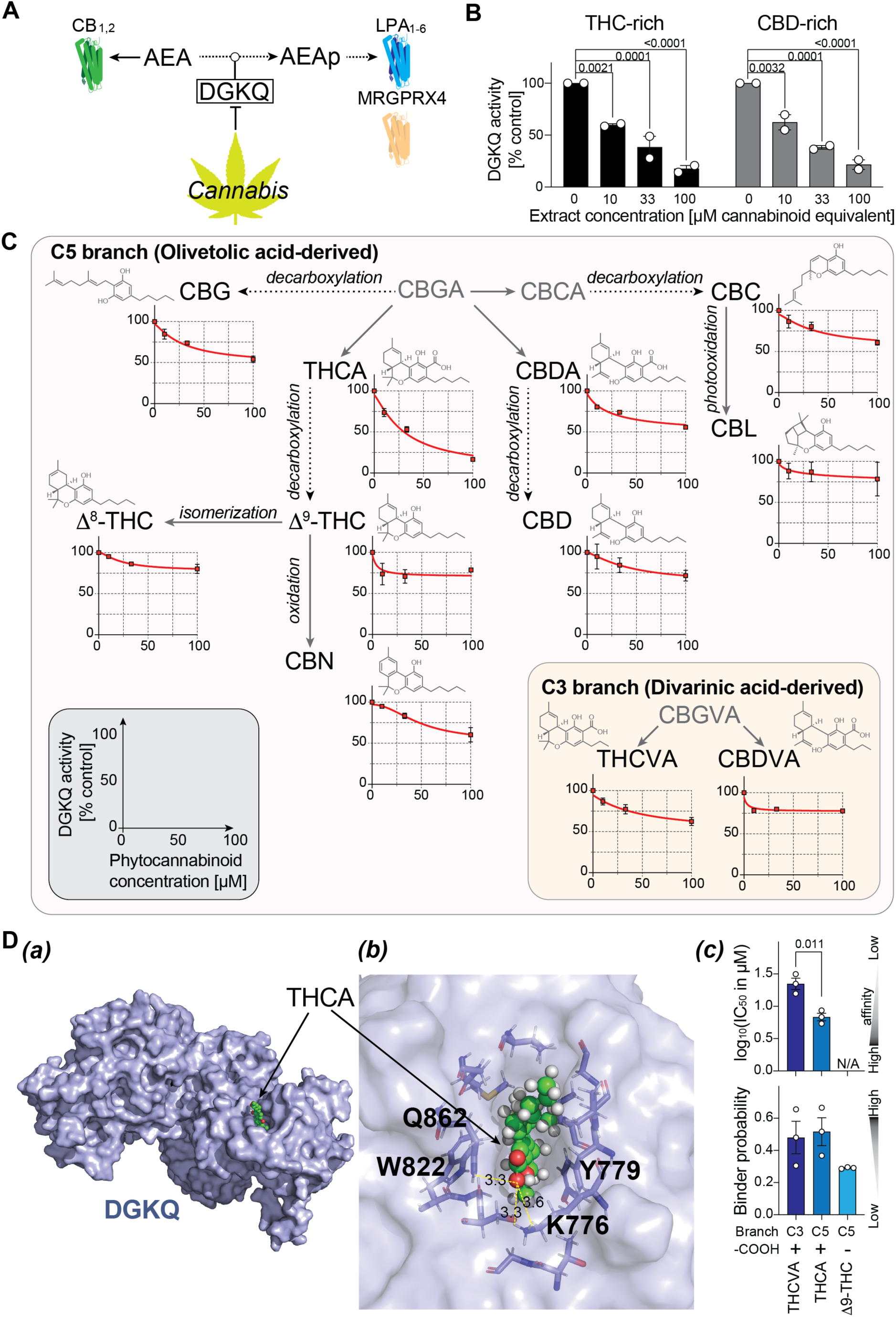
Phytocannabinoids inhibit DGKQ to modulate the AEA-to-AEAp signaling switch. **(A)** Conceptual model showing DGKQ as an enzymatic switch that phosphorylates AEA to generate AEAp. AEA engages canonical cannabinoid receptors CB_1/2_, whereas AEAp favors lysophospholipid-type GPCR signaling through LPARs and MRGPRX4. Phytocannabinoids are proposed to inhibit DGKQ activity, thereby affecting AEA-to-AEAp conversion. **(B)** THC-rich and CBD-rich *Cannabis* extracts inhibit AEAp production by DGKQ. Recombinant human DGKQ was incubated with AEA and ATP in the presence of increasing concentrations of *Cannabis* extracts, expressed as µM cannabinoid equivalent, and activity was measured using the ADP-Glo^TM^ assay. DGKQ activity is shown relative to vehicle control. Individual data points and summary bars are shown; P values, two-way ANOVA followed by Tukey’s multiple comparisons test. **(C)** Mapping DGKQ inhibition onto phytocannabinoid biosynthetic relationships. C5 cannabinoids derived from the olivetolic acid branch and C3/varin cannabinoids derived from the divarinic acid branch are shown together with their chemical relationships. Plots show DGKQ activity relative to vehicle control across phytocannabinoid concentrations. Red lines serve as visual guides. Gray labels denote biosynthetic precursors or intermediates not directly tested in the DGKQ inhibition assay. DGKQ inhibition was distributed across multiple phytocannabinoid branches, with the strongest observed inhibition by tetrahydrocannabinolic acid (THCA), moderate observed inhibition by cannabigerol (CBG), cannabidiolic acid (CBDA), cannabinol (CBN), cannabichromene (CBC), and tetrahydrocannabivarinic acid (THCVA), and weaker observed inhibition by cannabidiol (CBD), cannabidivarinic acid (CBDVA), cannabicyclol (CBL), Δ^9^-tetrahydrocannabinol (Δ^9^-THC), and Δ^8^-THC. **(D)** Predicted binding of THC-type phytocannabinoids to DGKQ. ***(a)*** Structural modeling placed THCA within the putative DGKQ substrate-binding pocket. ***(b)*** Enlarged view showing the THCA carboxyl group positioned near Q862 and K776, with predicted polar contacts indicated by dashed lines and distances shown in Å. The aromatic ring is positioned between Y779 and W822, suggesting additional hydrophobic and/or aromatic interactions within the pocket. Together, these predicted contacts provide a possible structural rationale for the strong DGKQ inhibition observed with THCA. ***(c)*** Boltz-2 affinity scoring predicted a significantly lower log_10_[IC_50_ in µM] for THCA than the C3 analog THCVA (p = 0.011, unpaired t-test, mean ± SEM, n = 3 independent prediction). No affinity value was reported for Δ^9^-THC (N/A), because its predicted binder probability was low (< 0.30; mean ± SEM, n = 3 independent prediction).

Because *Cannabis* extracts comprise a heterogenous mixture of structurally related, but distinct, molecules, we screened 11 extract constituents for inhibitory activity using recombinant DGKQ. Mapping these activities onto phytocannabinoid biosynthetic relationships showed that DGKQ inhibition was distributed across multiple phytocannabinoid scaffolds (**Fig. 4C**). Among the tested compounds, tetrahydrocannabinolic acid (**THCA**) showed the strongest inhibition (∼80%), whereas its decarboxylated counterpart Δ^9^-THC and the related isomer Δ^8^-THC showed weaker inhibition (∼20%). A similar trend was observed between cannabidiolic acid (**CBDA**) and its decarboxylated counterpart CBD, suggesting a role for the carboxylic acid group in explaining the structural basis of DGKQ inhibition. In addition, tetrahydrocannabivarinic acid (**THCVA**) and cannabidivarinic acid (**CBDVA**), which contain shorter C3 alkyl side chains, showed weaker inhibition than their C5 counterparts THCA and CBDA, suggesting that the pentyl side chain may contribute to DGKQ inhibition. Among the remaining non-acidic phytocannabinoids, cannabigerol (**CBG**), cannabichromene (**CBC**), and cannabinol (**CBN**) inhibited DGKQ more effectively (∼50%) than cannabicyclol (**CBL**) (∼20%). To explore the structural basis of phytocannabinoid inhibition, we modeled THCA, THCVA, and Δ^9^-THC binding to DGKQ (**Fig. 4D**). THCA was predicted to bind within the DGKQ substrate-binding pocket, with modeling favoring THCA over THCVA and Δ^9^-THC, likely due to favorable interactions with its carboxylic acid and longer hydrophobic alkyl chain (**Fig. 4D**). Collectively, these activity patterns suggest emerging structure-activity trends. DGKQ inhibition is favored by an acidic THC-type scaffold and a C5 alkyl chain, whereas decarboxylation, varin-type C3 side chains, or less favorable ring geometries reduced inhibitory activity.

## Discussion

In this study, we identified DGKQ as a biochemical link between the eCB and LPA signaling systems, showing that eCBs have greater signaling flexibility than previously appreciated. In addition to functioning as ligands for CB_1_ and CB_2_, eCBs are also branch-point metabolites that can be phosphorylated to alter signaling output. In this model, DGKQ converts AEA into AEAp (**Figs. 2 and 3**), which engages LPARs and MRGPRX4, and converts 2-AG into 2-arachidonoyl-LPA, an LPAR agonist (**Figs. 2 and 3**). DGKQ therefore establishes a direct enzymatic connection between two previously distinct lipid signaling systems that had mainly been linked through genetic, physiological, or compensatory observations (*34-36*).

The established signaling activity of 2-arachidonoyl-LPA through LPARs, together with the AEAp-LPARs/MRGPRX4 pathways identified here, defines an emerging class of lipid mediators, phospho-eCBs. A potential physiological role for phospho-eCBs is suggested at synapses in the central nervous system: 2-AG is a well-established retrograde messenger that travels from postsynaptic to presynaptic terminals, where it activates CB_1_ to suppress neurotransmitter release (*6, 7, 37*). Recent studies further show that 2-AG can be released and transported in extracellular microvesicles (*38*), providing a potential mechanism for its delivery across the synaptic space. In addition to activating CB_1_, 2-AG arriving at the presynaptic terminal can be metabolized by monoacylglycerol lipase to terminate eCB signaling (*39*). Our findings raise the possibility of an additional, functionally opposing pathway. DGKQ localizes to excitatory presynaptic terminals (*40*), where its catalytic activity is required for efficient synaptic-vesicle recycling (*40*). In addition, DGKQ could phosphorylate incoming 2-AG to generate 2-arachidonoyl-LPA. This product could, in turn, engage presynaptic LPA_2_ signaling, which is known to increase glutamate-release probability (*41, 42*). Notably, 2-arachidonoyl-LPA was the most potent LPA species at LPA_2_ in our assays (**Fig. 1B**), supporting the possibility that DGKQ-mediated phosphorylation could convert an inhibitory retrograde eCB signal into a facilitatory signal at the synapse.

Another potential physiological role for the phospho-eCB AEAp is suggested in skin biology, based on the expression patterns of DGKQ, CB_1_, LPA_1_/LPA_5_ and MRGPRX4 and their reported functions. A phosphorylation-dependent reversal analogous to the 2-AG/2-arachidonoyl-LPA axis may occur in keratinocytes, where DGKQ expression is supported by multiple single-cell RNA-seq datasets (*43*). AEA-CB_1_ signaling suppresses keratinocyte differentiation (*44*). Conversely, LPA_1_ and LPA_5_, among the LPARs most responsive to AEAp in our assays (**Fig. 1B**), are activated by AEAp with potencies approaching those of 2-arachidonoyl-LPA and promote keratinocyte differentiation and epidermal barrier formation (*45*). Thus, DGKQ-mediated conversion of AEA to AEAp could redirect an inhibitory AEA signal toward differentiation-promoting LPAR pathways. Phosphorylation of 2-AG to 2-arachidonoyl-LPA could potentially reinforce the same LPAR output. Moreover, MRGPRX4 is expressed in itch-sensing human sensory neurons and mediates bile acid-evoked cholestatic itch (*16, 17*). Because AEAp activates MRGPRX4, locally generated AEAp could provide an additional route through which DGKQ-dependent eCB phosphorylation modulates cutaneous sensory signaling. Together, these examples suggest that phosphorylation can act as a molecular switch that redirects eCB signaling toward distinct GPCR pathways and physiological outputs.

The inhibition of DGKQ by phytocannabinoids (**Fig. 4**) further suggests that this enzymatic switch can be pharmacologically tuned. This inhibition could shift AEA metabolism away from AEAp production and may alter the balance between canonical eCBs and LPAR/MRGPRX4-active phospho-eCBs. Structural modeling reveals that inhibition depends on scaffold fit, with two key features involving polar contacts to the carboxylic acid on acidic phytocannabinoids, and hydrophobic complementarity with the alkyl chain. This interaction with DGKQ provides a potential chemical basis for *Cannabis* bioactivities that have not yet been mechanistically explained. In addition, the convergence of multiple phytocannabinoid scaffolds on DGKQ positions this enzyme as one candidate molecular node for modulation of eCB metabolism beyond direct CB_1_/CB_2_ activation. For example, at excitatory synapses, inhibition of DGKQ could reduce formation of the LPA_2_-active phospho-eCB, thereby weakening a facilitatory pathway that opposes CB_1_-mediated suppression of neurotransmitter release. In theory, phytocannabinoid inhibition of DGKQ could enhance the net inhibitory effect of cannabinoid signaling without necessarily increasing CB_1_ activity itself. A similar mechanism may operate in cutaneous sensation. *Cannabis* and cannabinoid-based interventions have been reported to reduce pruritus in humans (*46*), although the underlying mechanisms are not well established. Because AEAp activates the human itch receptor MRGPRX4, inhibition of DGKQ could reduce local AEAp production and thereby provide an additional mechanism for the antipruritic effects of phytocannabinoids alongside canonical cannabinoid receptor signaling. Whether this pathway operates *in vivo* warrants further investigation.

More broadly, the clinical relevance of the AEA-DGKQ-AEAp-LPAR/MRGPRX4 pathway is suggested by genetic and functional links between DGKQ and diseases. Variants at the DGKQ locus have recently been associated with Parkinson’s disease risk (*47-52*), and altered DGKQ expression was found in several human neurodegenerative disease datasets (*53, 54*). DGKQ is also a susceptibility locus for systemic sclerosis, with associated variants suggesting regulatory effects on DGKQ expression (*55, 56*). Together with the broad involvement of eCB and LPA signaling in neurological, inflammatory, and sensory processes (*1, 6, 12*), these observations suggest that DGKQ may represent a disease-relevant node controlling the balance between canonical eCB and phospho-eCB signaling. Determining whether modulation of this pathway contributes directly to disease phenotypes, and whether it can be therapeutically targeted, will require further study.

## Acknowledgments

We thank Dr. A. Inoue for providing the TGF-α shedding assay, Richard Rivera for experimental support, and Danielle Jones for administrative support. The authors also acknowledge instrumentation resources at the UC San Diego Agilent Center of Excellence in Cellular Intelligence.

## Author contributions

Conceptualization: YK

Methodology: AL, YK

Investigation: AL, YK, HM, KN, DC, DJS, VT

Funding acquisition: BM, JC

Project administration: YK, BM, JC

Supervision: YK, BM, JC

Writing – original draft: AL, YK

Writing – review & editing: AL, YK, HM, KN, DC, DJS, VT, BM, JC

## Data and materials availability

All data are available in the main text or the Supplementary materials.

### STAR Methods

### General

This project was registered with and conducted under the oversight of the University of California, San Diego Controlled Substances Program, the Research Advisory Panel of California, and the US Drug Enforcement Agency (DEA). All researchers who accessed and performed work with controlled substances passed background screening. All experiments using controlled substances were performed in a DEA-registered laboratory, and all protocols using controlled substances were reviewed by the relevant authorities, including product acquisition, inhibition assays, and product disposal. To prevent illicit use, compounds and used plates containing controlled substance residues were stored in a locked -20°C freezer fixed to the benchtop using Guardianite heavy-duty safety lock pads and Master four-digit combination locks. The freezer was similarly secured shut using the same safety lock pads and Master four-digit combination lock. The logbook was secured in a biometric safe accessible only to individuals registered with the UC San Diego Controlled Substances Office and approved following background check to work with controlled substances.

### GPCR assays (intracellular Ca^2+^ signaling, backscattering interferometry-based binding assay, and TGF-α shedding assay)

Human LPAR1-6 and MRGPRX4 expression constructs were generated in pCXN2.1 vectors containing an N-terminal hemagglutinin (HA) epitope tag. Plasmids were transfected into B103, CHO, HEK293, or Neuro2a cells using Lipofectamine 2000 (Thermo Scientific, Cat# 11668030). Stable cell lines were generated by selection with G418 (1 mg/ml, Thermo Scientific, Cat# 10131035) for 2 weeks followed by sorting of HA-positive populations using a FACSAria II cell sorter (BD Biosciences: RRID:SCR_018934). Cells were maintained in DMEM high glucose supplemented with 10% fetal bovine serum (Corning, Cat# 35011CV) and penicillin-streptomycin (Thermo Scientific, Cat# 15140122). Cell-surface receptor expression was confirmed by flow cytometry using an anti-HA antibody (clone 3F10, Roche: RRID:AB_390918) (*1-3*).

For intracellular Ca^2+^ mobilization assays, cells were plated in black clear-bottom 384-well plates (20,000 cells/well) and serum-starved in FreeStyle^TM^ 293 Expression Medium (Thermo Scientific, Cat# 12338018) for at least 3 h prior to assay. For GPCR screening experiments, cells were transiently transfected with untagged GPCR expression constructs in pcDNA3.1 on the day of plating. Cells were loaded with Fluo-4 dye (FLIPR Calcium 4 Assay Kit, Molecular Devices, Cat# R8143) according to the manufacturer’s instructions, and fluorescence changes were monitored using an FDSS7000 (Hamamatsu) with excitation at 485 nm and emission at 525 nm (*2*).

Backscattering interferometry (BSI) binding assays were performed as previously described (*2*). Briefly, membrane nanovesicles prepared from receptor-expressing or control cells (20 µg/ml total protein) were incubated with lipid ligands for 1 h at room temperature and loaded into microfluidic channels for analysis. Specific binding signals were calculated by subtracting the control nanovesicle refractive index (non-specific binding) from receptor-containing nanovesicle refractive index (total binding). Binding responses against AEAp (MilliporeSigma, Cat# 870440C) or LPA (Avanti, Cat# A85130) were expressed as changes in milliradians (Δmrad) and analyzed by nonlinear regression using Prism (GraphPad: RRID:SCR_002798) (*2*).

TGF-α shedding assays were performed as previously described (*4*). HEK293 or Neuro2a cells were seeded onto 6-well plates one day prior to transfection. Cells were co-transfected with plasmids encoding AP-TGF-α and the MRGPRX4 constructs. Twenty-four hours after transfection, cells were stimulated with ligands for DCA (Sigma-Aldrich, Cat# D2510) and AEAp (MilliporeSigma, Cat# 870440C) at the indicated concentrations. Alkaline phosphatase activity released into conditioned medium and that remaining in the cells was quantified using p-nitrophenyl phosphate (MilliporeSigma, Cat# 4876), and GPCR activation was calculated as the percentage of AP-TGF-α released relative to total AP activity (*4*).

### Determination of AEA kinase activity in tissue homogenates

All animal protocols were approved by the Institutional Animal Care and Use Committee (IACUC) of the Sanford Burnham Prebys Medical Discovery Institute and conform to National Institutes of Health guidelines and public law. Tissues were collected from C57BL/6 mice, flash frozen, and stored at -80°C until use. Tissues were weighed and homogenization buffer containing 50 mM HEPES (pH 7.5), 300 mM sucrose, and 1x cOmplete Protease inhibitor cocktail (Millipore Sigma, Cat# 11873580001) was added for a final concentration of ∼10% w/v tissue to buffer. Tissues were homogenized until the mixture appeared homogenous and no chunks remained. Homogenate was then centrifuged at 10,000×*g* for 10 min at 4°C, and the supernatant was either collected and flash-frozen in liquid nitrogen or carried on for further clarification. The 10,000×*g* supernatant was further centrifuged at 20,000×*g* for 10 min at 4°C, and the pellet was dissolved in homogenization buffer. Both supernatant and dissolved pellet were flash frozen on liquid nitrogen. All tissue homogenates and resuspended pellets were analyzed using a Bradford assay (Bio-Rad, Cat# 5000205) to determine total protein concentration prior to flash freezing in liquid nitrogen. All tissue homogenates and resuspended pellets were stored at -80°C until use.

Mouse brain and liver homogenates (10,000×*g* supernatants) were incubated at 1 mg/mL final concentration in Eppendorf tubes (1.7 mL) containing 200 µM AEA (Avanti, Cat# A87430) or AEA-d4 (Avanti, Cat# A85462), 2 mM ATP (MilliporeSigma, Cat# A2383), 1 mM Na_3_VO_4_ (Millipore Sigma, Cat# 450243) 30 mM MgCl_2_ (MilliporeSigma, Cat# M8266) and 1x phosphatase inhibitor cocktail (Thermo Scientific, Cat# AAJ63907AA), in 50 mM HEPES buffer (pH 7.5) for 30 min at 37°C in a final reaction volume of 100 µL. To stop reactions, 100 uL of methanol was added to each reaction, and samples were briefly vortexed. Precipitated proteins were pelleted by centrifugation at 15,000×*g* for 5 min, and supernatants were passed through 0.2 µm PTFE filters by centrifugation at 4,000×*g* before LC-MS analysis. Filtered samples (5-30 µL injection volume) were analyzed using an Agilent Technologies 1260 Infinity series HPLC equipped with a degasser, binary pump, autosampler, and diode array detector coupled to a 6530 Accurate-Mass Q-TOF MS equipped with a Kinetex 5 μm C18 100 Å, 150 × 4.6 mm column. The solvents used were mass spectrometry grade, and solvent A was Optima^TM^ LC-MS grade water (Thermo Scientific, Cat# W6-1) + 0.1% Optima^TM^ LC-MS grade formic acid (FA, Thermo Scientific, Cat# A117-50), while solvent B was Optima^TM^ LC-MS grade acetonitrile (Thermo Scientific, Cat# A955-1) + 0.1% FA. Analysis was performed at a 0.750 mL/min flow rate with equilibration at 50% solvent B for 2 min, followed by 8 min linear gradient to 100% solvent B, and held at 100% solvent B until 17 min, followed by a linear 1 min gradient back to 50% solvent B, and held there for 2 min to end the method. Data analysis was performed using MassHunter Workstation software version B.05.01 and traces were exported to Microsoft Excel and GraphPad Prism for processing.

For heat treated reactions, tissue homogenates were diluted to 1 mg/mL and heated at 95°C for 5 min before the other reaction components were added. For assays without ATP, ATP was withheld from the reaction mixture. To test the effects of divalent cations, other cations were added at 10 mM either in addition to or in place of MgCl_2_. To test the effects of 2-AG (Avanti, Cat# A87450) on AEAp production, 2-AG was additionally added to the reaction mixture in an equimolar amount (200 µM).

### Activity-guided fractionation, proteomics analysis, and inhibitor screening

Brain and liver homogenates were prepared from pooled tissues collected from male mice as described above. Homogenate was prepared as described above and centrifuged at 10,000×*g* for 10 min. Supernatant protein concentrations were measured using a Bradford assay (Bio-Rad, Cat# 5000205), and homogenization buffer was added to normalize protein concentration of brain and liver samples to 3.5 mg/mL. Tissue homogenates (2 mL each) were loaded onto a size-exclusion column containing Sephadex S200 equipped to a fast protein liquid chromatography Cytiva AKTA Pure 25 L1 system fitting with a F9-C fraction collector and an S9 sample pump and controlled by Unicorn v.7 software. Proteins were eluted using a flow rate of 1.0 mL/min with 50 mM HEPES buffer (pH 7.5) containing 100 mM NaCl, for 120 min. PMSF (1 mM, Millipore Sigma, Cat# P7626) was added to each fraction (5 mL each), and fractions stored at 4°C overnight. Fractions then were concentrated to at least 2 mg/mL and used in assay conditions described above to test for AEAp production.

Brain and liver fractions (three each; six samples total) were submitted to the UCSD Proteomics Core Facility for analysis using an Orbi-Trap Fusion LUMOS instrument equipped with a C18 column. Data analysis was performed using PEAKS Studio 12 (Bioinformatics Solutions) with a false discovery rate of 1%, and the data were compiled in Excel. All proteins containing “kinase” were compiled and filtered; only kinases with >10% coverage in “brain fraction 2” and <10% coverage in *all* other fractions were considered. From this pool, eleven candidates known act on small molecules or metabolite substrates were selected.

Whole brain homogenate (10,000×*g* supernatant) was assayed for AEAp production as described above in the presence of small molecule-inhibitors at 100 µM, including metformin, BAMB4 (Cayman Chemicals, Cat# 36583), R-59-022 (Cayman Chemicals, Cat# 16772), PANKi (Cayman Chemicals, Cat# 31002), and SKII (Cayman Chemicals, Cat# 10009222). Samples were run in triplicate and normalized to control samples where no inhibitor was added.

### LC-MS/MS-based AEA kinase assay using recombinant DGKQ

Recombinant human DGKQ was purchased from Carna Bioscience® (Cat# 12-109) and used without further purification. Recombinant enzyme assays were performed using 50 ng of protein in 50 mM HEPES buffer (pH 7.5) containing 30 mM MgCl_2_, 2 mM ATP, and 200 µM AEA (Avanti, Cat# A87430) or AEA-d4 (Avanti, Cat# A85462). Reactions were incubated at 37°C for 30 min, quenched with 100 µL MeOH, centrifuged, filtered, and analyzed by mass spectrometry as described above. To improve the signal to noise ratio and assay sensitivity, metal, pH, and nucleoside triphosphate donor preferences were characterized using an Agilent Technologies 1290 Infinity II series UPLC equipped with a degasser, binary pump, autosampler, and diode array detector coupled to a 6495D triple quadrupole (QQQ) mass spectrometry equipped with an Agilent Poroshell 120 2.7 µm EC-C18 2.1 x 50 mm column. The solvents used were mass spectrometry grade, and solvent A was Optima^TM^ LC-MS grade water (Thermo Scientific, Cat# W6-1) + 0.1% Optima^TM^ LC-MS grade FA (Thermo Scientific, Cat# A117-50), + 5 µM Agilent InfinityLab Deactivator Additive (Agilent Technologies, Cat# 5191-4506), and solvent B was Optima^TM^ LC-MS grade acetonitrile + 0.1% FA (Thermo Scientific, Cat# A117-50). The method involved a 0.5 mL/min flow rate, equilibration at 60% solvent B for 0.5 min, followed by a linear 4 min gradient to 95% solvent B, and held at 95% solvent B for 1.5 min, followed by a 1 min post run to re-equilibrate back to 60% Solvent B. A multiple reaction monitoring (MRM) method was developed using AEAp (Cayman Chemicals, Cat# 10180) and 1-arachidonoyl-LPA (Cayman Chemicals, Cat# 10019) standards before analyzing reactions. Optimized collision energies for AEAp (34 V) and 1-arachidonoyl-LPA (71 V) were used with parent masses of 426.2 and 457.2 amu, respectively, and fragmentation mass of 78.9 (corresponding to loss of phosphate).

To determine DGKQ preferences for 1) metal ions, 2) pH, and 3) nucleoside triphosphate donors, assays were performed using the triple-quadrupole mass spectrometry method described above were carried out. For cation preferences, assays were performed using 50 ng of DGKQ per assay in 50 mM HEPES buffer (pH 7.5) containing 2 mM ATP, 200 µM AEA, and either 30 mM MgCl_2_, or 10 mM of either ZnCl_2_ (MilliporeSigma, Cat# 208086) or MnCl_2_ (MilliporeSigma, Cat# 328146). For pH tests, assays were performed using 50 ng of DGKQ per assay in 50 mM HEPES buffer at pH values of either 5.5, 6.5, 7.5, 8.5, or 9.5, containing 30 mM MgCl_2_, 200 µM AEA, and 2 mM ATP. For nucleoside triphosphate preferences, 50 ng of DGKQ was added per assay in 50 mM HEPES buffer (pH 7.5) containing 30 mM MgCl_2_, 200 µM AEA, and either ATP (2 mM), GTP (2 mM, Millipore Sigma, Cat# G8877) or both. Assays were kept at 37°C for 30 min, before quenching with 100 µL MeOH followed by centrifugation at 15,000×*g* to pellet precipitated protein. Supernatant was filtered using 0.2 µm PTFE filters by centrifugation at 4,000×*g* prior to LC-MS analysis using the QQQ mass spectrometer and method described above with an injection volume of 5 µL. Data analysis was performed using OpenLab CDS and MassHunter workstation software, and LC traces were exported to Excel and GraphPad Prism software for processing.

### Screening eCB kinase activity across DGK isozymes

Individual plasmids, each encoding a different DGK family member and eGFP under separate promoters were obtained from VectorBuilder and transfected into HEK293 cells using Lipofectamine 3000. Cells were pelleted and flash frozen at -80°C prior to analysis. Cell pellets were resuspended in 1 mL of 50 mM HEPES buffer (pH 7.5) and sonicated at 40% amplitude for 2 s on and 2 s off five times to lyse. Cell lysates were clarified at 15,000×*g* at 4°C for 5 min to pellet debris. Supernatants were analyzed using Bradford assay (Bio-Rad, Cat# 5000205) to determine total protein concentration. Reactions were performed using 0.5 mg/mL final protein concentration, 2 mM ATP, 1 mM Na_3_VO_4_, 1x phosphatase inhibitor cocktail (Thermo Scientific, Cat# AAJ63907AA), 30 mM MgCl_2_, and 200 µM AEA or 2-AG, at 37°C for 30 min before being quenched with 100 µL MeOH. Samples were centrifuged at 15,000×*g* for 10 min to pellet precipitated proteins. The supernatant was passed through a 0.2 µm PTFE filter, and the flow through was transferred to a mass spectrometer vial for analysis. Samples were analyzed using the QQQ method described above with an injection volume of 10 µL. MRM traces were exported to Excel, and the chromatographic peak areas were extracted to generate the graphs shown in Figure 3A and B.

### ADP-Glo^TM^-based eCB kinase assay using recombinant DGKQ

A commercial ADP-Glo^TM^ Assay kit (Promega®, Cat# V9101) was used according to the manufacturer’s instructions to determine biochemical parameters of DGKQ (Carna Bioscience®, Cat# 12-109) with AEA (Avanti, Cat# A87430) and 2-AG (Avanti, Cat# A87450) Assays were performed in 384-well white plates (Grenier Bio-One®). Assay wells were separated by one empty well to minimize luminescence bleed over from wells with high signal intensity. Briefly, kinase reactions were performed in 5 µL reaction volumes containing 50 mM HEPES (pH 7.5), 10 mM MgCl_2_, 1 mM ATP, 50 ng DGKQ, and substrates. Reactions were initiated by the addition of DGKQ and incubated for 30 min at room temperature (∼22°C). Following incubation, 5 µL ADP-Glo™ Reagent was added to each well to terminate the kinase reaction and deplete remaining ATP. Plates were incubated for 40 min at room temperature. Subsequently, 10 µL Kinase Detection Reagent was added to convert ADP to ATP and generate a luminescent signal. Following an incubation of 60 min at room temperature, luminescence was measured using a Tecan SPARK^TM^ plate reader with a 1,000 ms integration time. Luminescence values were proportional to ADP production and used as a measure of kinase activity. Background signal was determined using no-enzyme control reactions and subtracted from all samples. Kinetic data obtained at 0.5 mM and 1.0 mM ATP were analyzed by global nonlinear regression using Python and the lmfit package. Mean luminescence values from replicate measurements were fitted simultaneously to either a single Michaelis-Menten model or a biphasic piecewise Michaelis-Menten model. For the single-phase model, each ATP condition was assigned an independent background and Vmax, while Km was shared across both ATP concentrations. For the biphasic model, the substrate response was modeled as two Michaelis–Menten components separated at 100 µM substrates. Background and phase-specific Vmax values were fitted independently for each ATP condition, whereas the apparent Km values for the low- and high-concentration phases were shared globally. Model performance was compared using Akaike information criterion (AIC). Fits were plotted together with mean ± SD values from replicate measurements.

To test the effects of phytocannabinoids, the protocol above was used with some modifications. Phytocannabinoid extracts, “THC-rich” and “CBD-rich” preparations, were acquired from the NIH NIDA Drug Supply Program. The extract was resuspended in dimethyl sulfoxide (Millipore Sigma, Cat# D8418). The dilutions for each extract were performed using the molecular weight of either Δ^9^-THC or CBD, the major constituent of each extract, to approximate a molar solution. Stock solutions of THCA (Cayman Chemicals, Cat# 33448), Δ^9^-THC (Cayman Chemicals, Cat#12068), Δ^8^-THC (NIDA Drug Supply Program, CAS: 5957-75-5), CBDA (Sanobiotec, Cat#G_SN38I), CBD (Millipore Sigma, Cat# C7515), THCVA (Cayman Chemicals, Cat #21259), CBDVA (Sanobiotec, Cat# G_SN47I), CBG (Millipore Sigma, Cat# SML4230), CBC (Sanobiotec, Cat# G_SN02I), CBN (Sanobiotech, Cat# G_SN03I), and CBL (Sanobiotech, Cat# G_SN48I), were likewise prepared in DMSO at 20 mM concentration. Phytocannabinoids were diluted in 50 mM HEPES (pH 7.5), and 1 µL of each diluted phytocannabinoid was added to a white 384-well plate (Grenier Bio-One®). No inhibitor wells contained 1 µL of only DMSO as a control. Next, 1.5 µL of DGKQ (50 ng total) was added to each well containing inhibitor, for a total of 2.5 µL of solution per well. The concentrations of each inhibitor per well after dilution with DGKQ was either 100 µM, 33 µM, or 10 µM. The plate was shaken for 30 s at 950 rpm to mix, spun down at 300×*g* for 2 min, and then incubated for 5 min at room temperature (∼22°C). Next, 2.5 µL of buffer mix containing 50 mM HEPES (pH 7.5), 20 mM MgCl_2_, 1 mM ATP, 400 µM AEA, and appropriate concentrations of phytocannabinoid inhibitor was added to each well such that the final concentrations of inhibitors were maintained at either 100 µM, 33 µM, or 10 µM. Final concentrations of MgCl_2_, ATP, and AEA in each reaction mixture were thus 10 mM, 500 µM, and 200 µM respectively. The plate was then shaken for 30 s at 950 rpm to mix, spun down at 300×*g* for 2 min, and then incubated for 30 min at room temperature. Following incubation, 5 µL of ADP-Glo™ Reagent was added to each well to terminate the kinase reaction and deplete remaining ATP. Plates were incubated for 40 min at room temperature. Subsequently, 10 µL Kinase Detection Reagent was added to convert ADP to ATP and generate a luminescent signal. After incubation for 60 min at room temperature, luminescence was measured using a Tecan SPARK^TM^ plate luminometer with 1000 ms dwell time. Luminescence values were proportional to ADP production and used as a measure of kinase activity. All inhibitor reactions were performed six times, three technical replicates across two biological replicates.

### Boltz-2 modeling

Boltz-2 protein-ligand modeling was performed using the Tamarind Bio web application with default settings (https://app.tamarind.bio/) (*5*). The DGKQ protein sequence (UniProt: P52824) was used as the protein input, and ligands were provided as SMILES strings. Predicted complex structures were inspected to assess ligand placement within the DGKQ substrate-binding pocket and proximity to candidate interacting residues. Boltz-2 affinity outputs were interpreted as model-derived scores rather than experimentally measured binding constants. Affinity scores are reported as log_10_[IC_50_ in µM], and binder ability is reported as binary binder probability. Three independent prediction runs were performed for each ligand, and values are presented as mean ± SEM.

## Key resources table

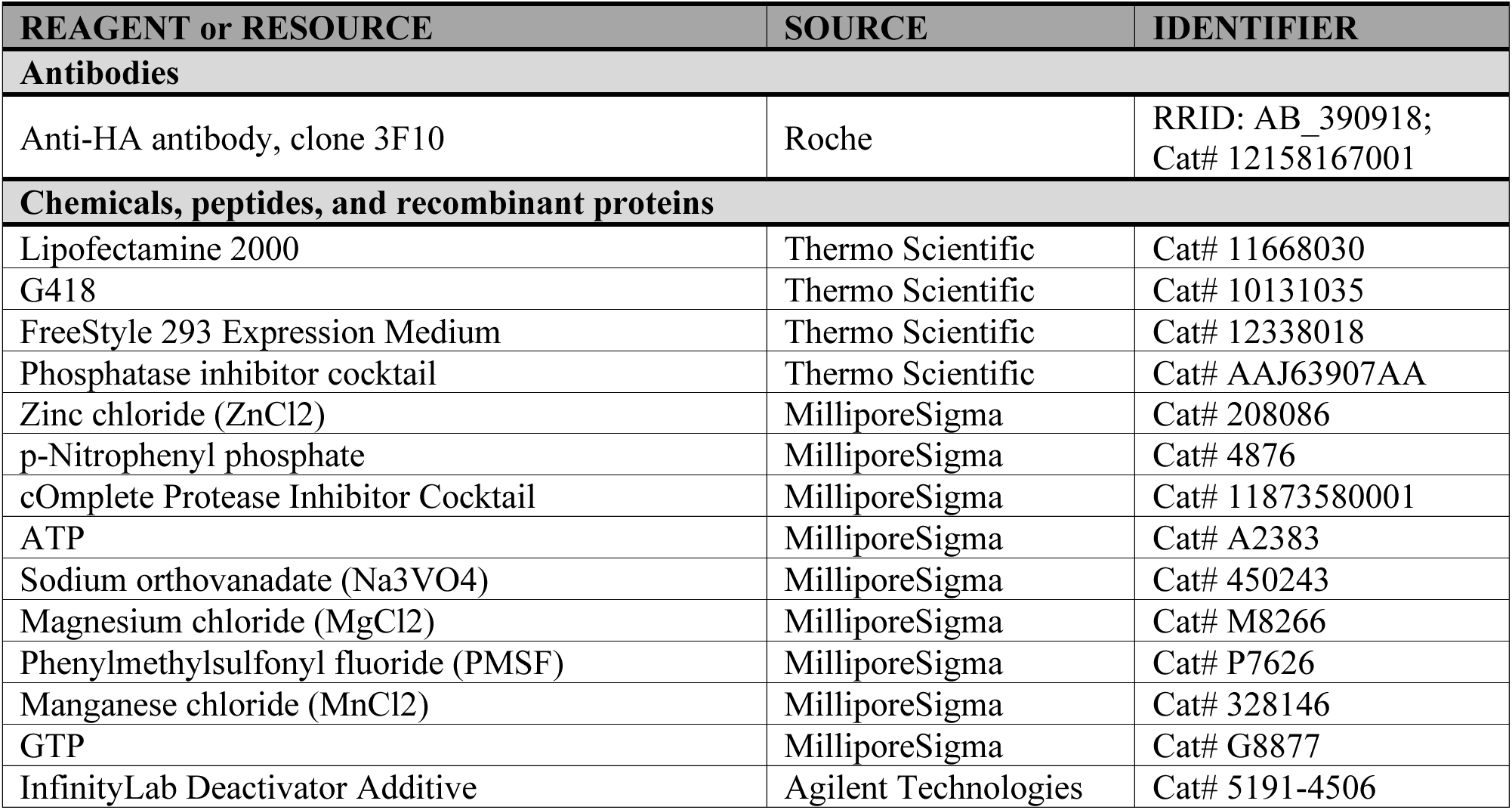
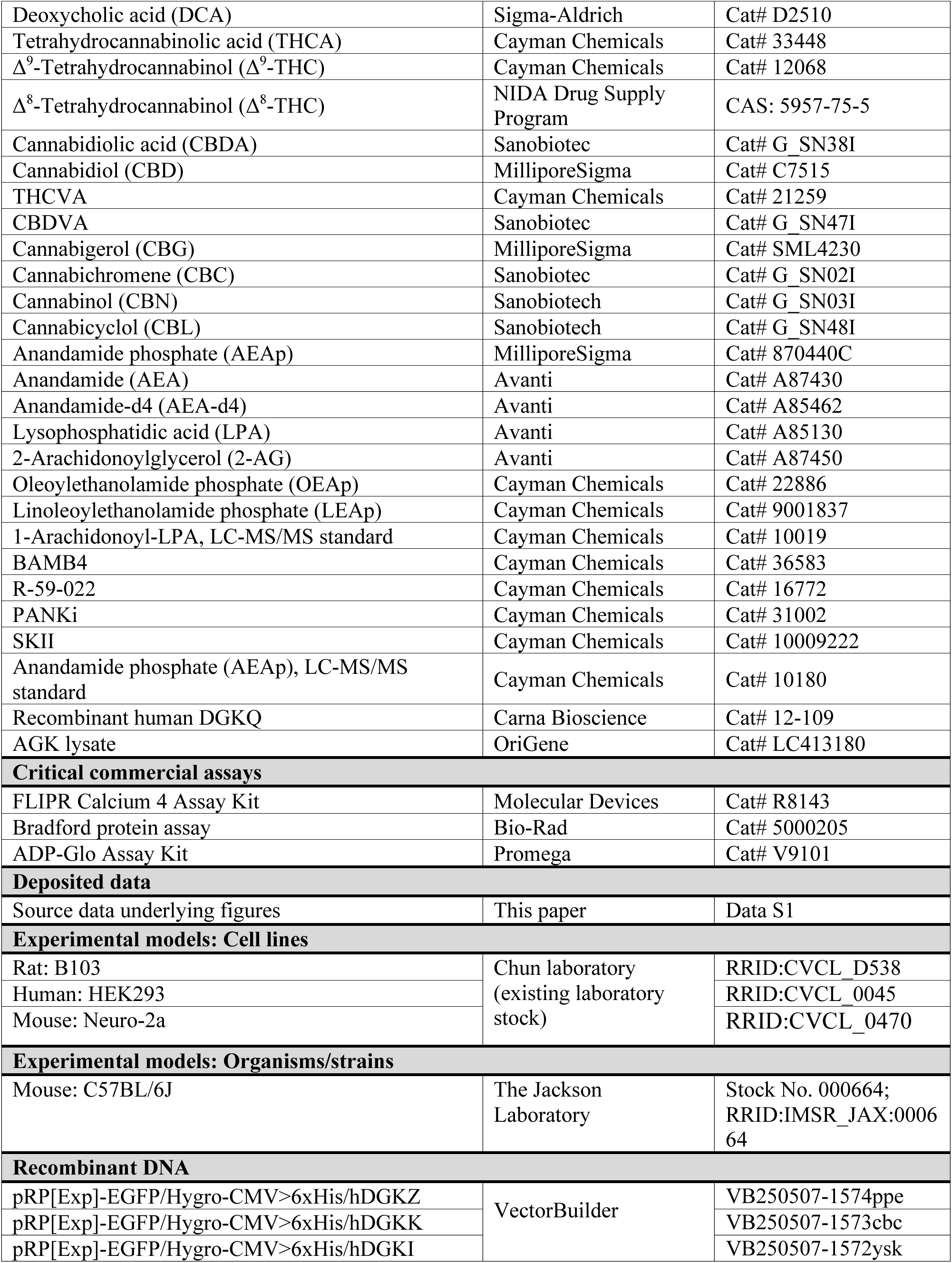
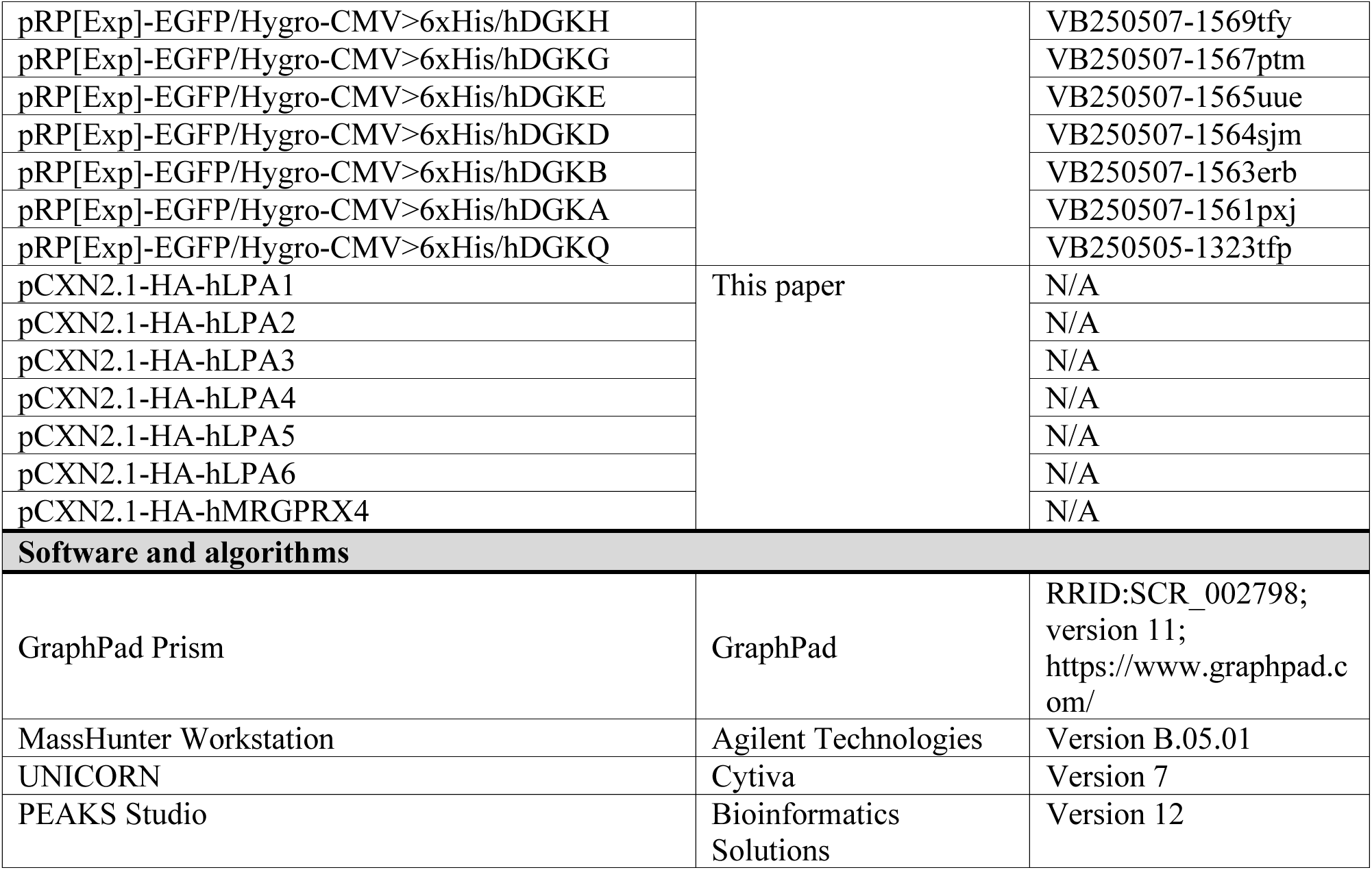

## Supplementary Figures for

**Fig. S1.**
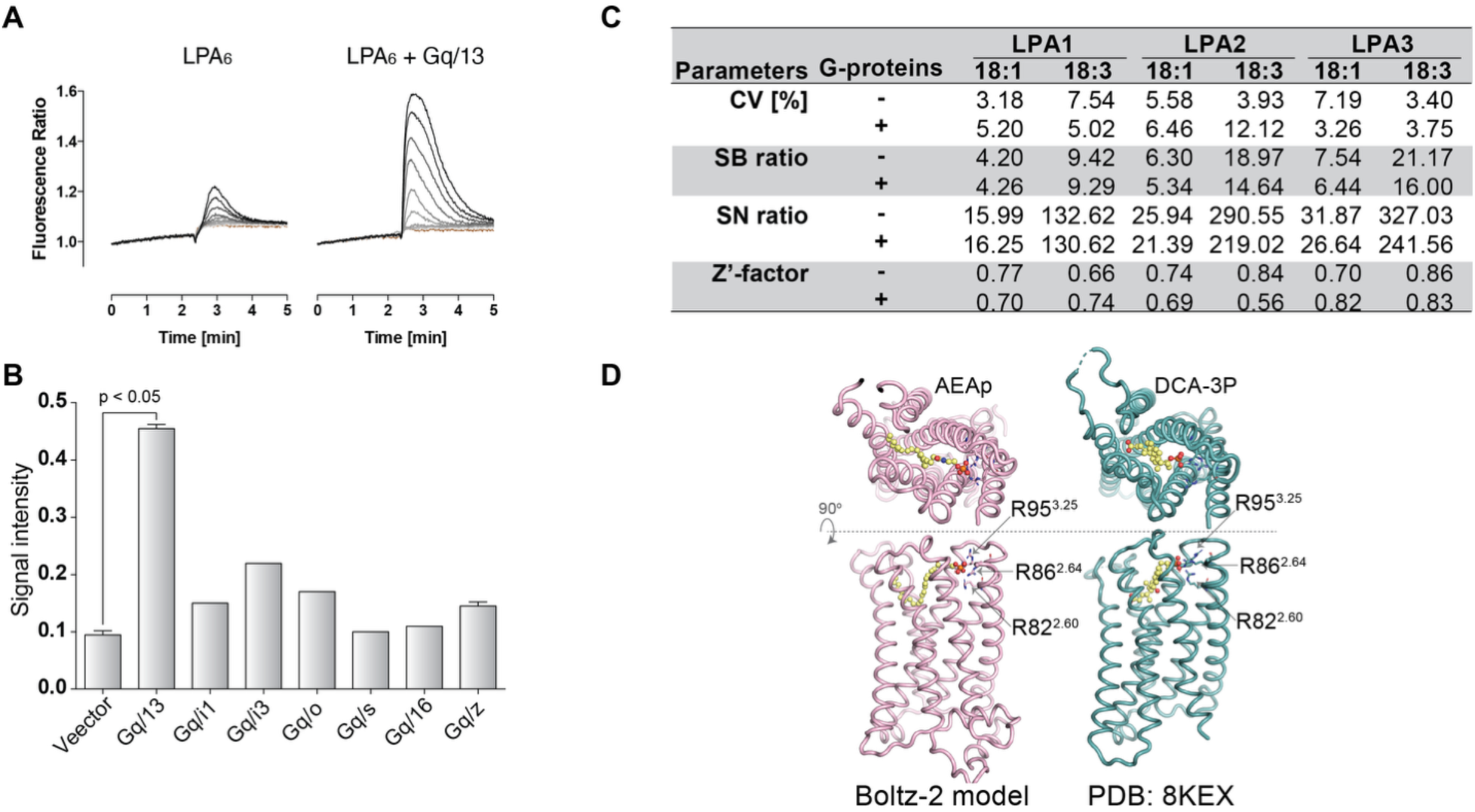
Optimization and validation of Ca^2+^ signaling assays used for LPA receptor and GPCR screening. **(A)** Representative intracellular Ca^2+^ traces in LPA_6_-expressing B103 cells stimulated with LPA in the absence or presence of a Gα_q_-Gα_13_-chimeric G protein. Co-expression of the chimeric G protein enabled to determine the LPA_6_-mediated Ca^2+^ mobilization. **(B)** Comparison of chimeric G proteins for coupling LPA_6_ activation to intracellular Ca^2+^ signaling. B103 cells expressing LPA_6_ were co-transfected with the indicated chimeric G proteins, and LPA-induced Ca^2+^ responses were quantified as signal intensity. The Gα_q_-Gα_13_-chimera produced the strongest response and was used for subsequent LPA_6_ assays. Data are mean ± SEM. P value was determined by one-way ANOVA. **(C)** Assay-quality metrics for representative LPA receptor Ca^2+^ assays. Coefficient of variation (CV), signal-to-background ratio (SB ratio), signal-to-noise ratio (SN ratio), and Z′-factor were calculated for LPA_1_, LPA_2_, and LPA_3_ assays stimulated with 18:1 or 18:3 LPA with or without chimeric G proteins. The resulting metrics supported the validity of these assay conditions for GPCR screening. **(D)** Structural comparison of the predicted AEAp-MRGPRX4 complex and the experimentally determined DCA-3P-MRGPRX4 complex. The AEAp-bound MRGPRX4 complex was modeled using Boltz-2 and compared with the cryo-EM structure of MRGPRX4 bound to phosphorylated deoxycholic acid (DCA-3P; PDB: 8KEX) (*6*). Both ligands position their phosphate groups near a positively charged arginine cluster, comprising R82^2.60^, R86^2.64^, and R95^3.25^, suggesting a shared phosphate-recognition mode.

**Fig. S2.**
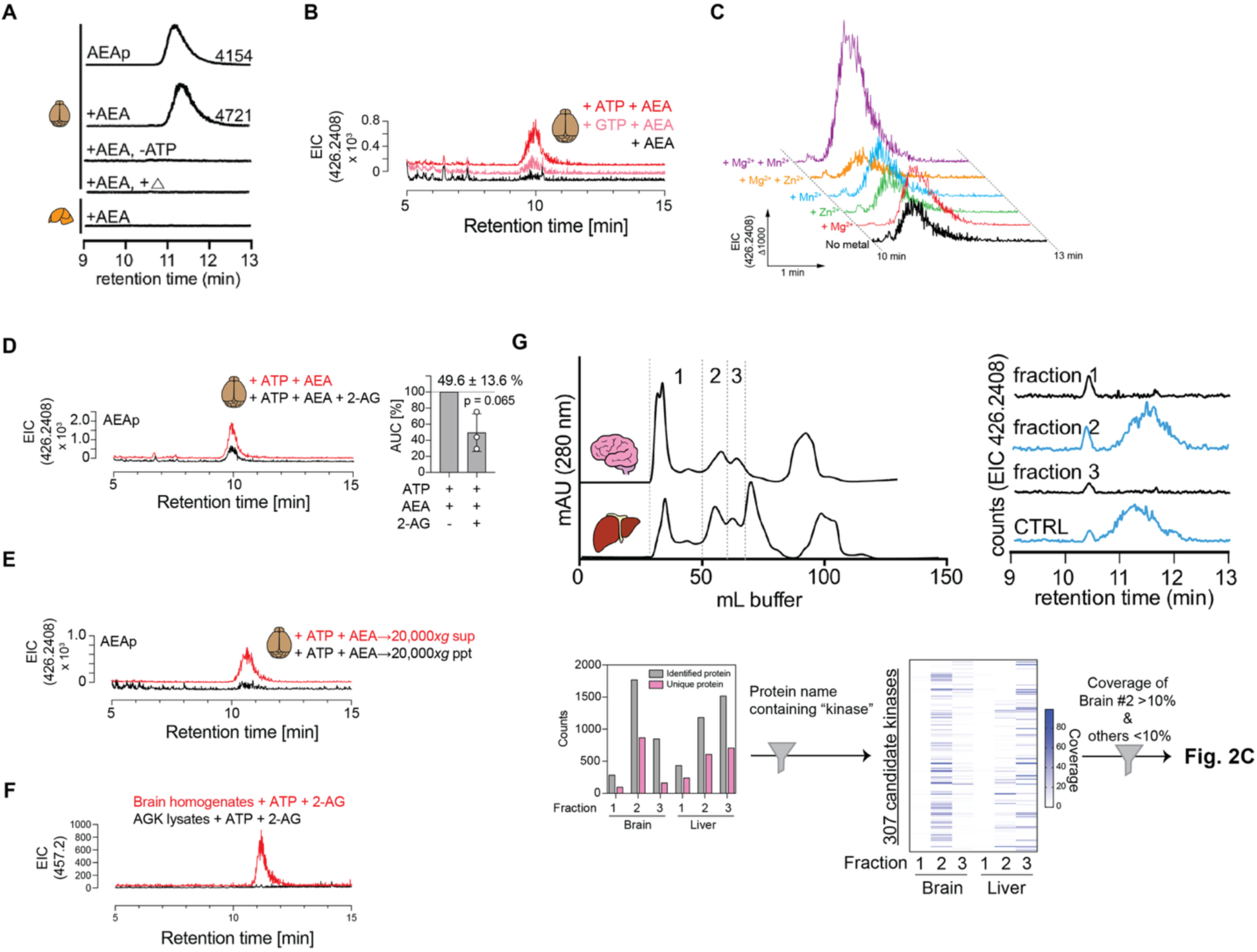
Biochemical characterization and activity-guided fractionation of brain AEA kinase activity. **(A)** LC-MS extracted ion chromatograms showing AEAp formation in tissue homogenate assays. An AEAp standard and mouse brain homogenates incubated with AEA and ATP showed peaks at the expected retention time for AEAp, whereas reactions lacking ATP, heat-inactivated brain homogenates, and liver homogenates showed little or no AEAp formation. **(B)** Nucleoside triphosphate donor preference of brain AEA kinase activity. Mouse brain homogenates were incubated with AEA in the presence of ATP or GTP, and AEAp formation was monitored by LC-MS. ATP supported AEAp formation more effectively than GTP under our assay condtions. **(C)** Effects of divalent cations on brain AEA kinase activity. Mouse brain homogenates were incubated with AEA and ATP in the absence or presence of Mg^2+^, Zn^2+^, Mn^2+^, or combinations of these cations. Mg^2+^ and Mn^2+^ increased AEAp formation, whereas Zn^2+^ did not enhance activity under these conditions. **(D)** Effect of 2-AG on AEA kinase activity in brain homogenates. Addition of 2-AG reduced AEAp formation, consistent with competition between AEA and 2-AG for a shared phosphorylation mechanism. Representative extracted ion chromatograms and quantification of AEAp peak area are shown. Data are mean ± SEM; P value is indicated. **(E)** Subcellular fractionation of brain AEA kinase activity. AEAp formation was detected primarily in the 20,000×*g* supernatant rather than the crude pellet fraction, arguing against enrichment of the activity in crude mitochondrial fractions. **(F)** 2-AG phosphorylation was observed in brain homogenates, but not in AGK lysates (OriGene, cat. #LC413180). **(G)** Activity-guided fractionation and proteomic prioritization of candidate kinases. Mouse brain and liver homogenates were separated by size-exclusion chromatography into three fractions. AEA kinase activity was detected only in brain fraction 2, as assessed by LC-MS detection of AEAp formation. Proteomic analysis identified proteins across brain and liver fractions, and proteins containing “kinase” in their name were prioritized. Candidate kinases were further filtered for >10% coverage in the active brain fraction and <10% coverage in other fractions, generating the candidate list shown in Fig. 2C.

**Fig. S3.**
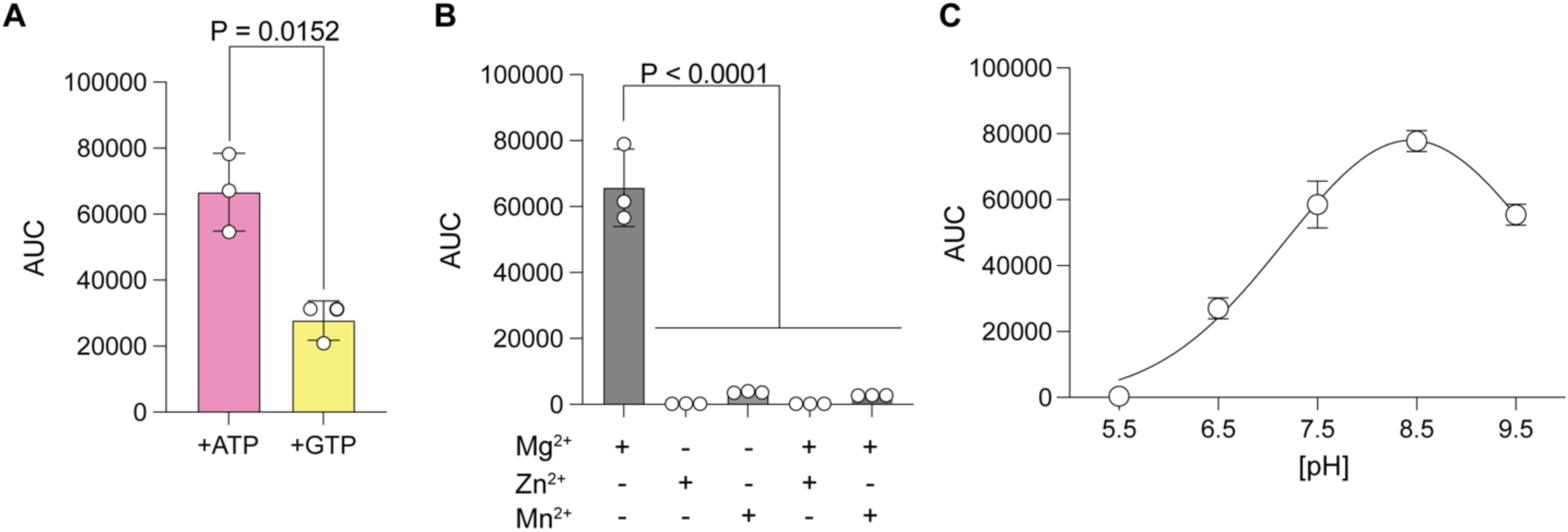
Biochemical characterization of recombinant DGKQ AEA kinase activity. **(A)** Nucleoside triphosphate donor preference of recombinant human DGKQ. DGKQ was incubated with AEA, Mg^2+^, and either ATP or GTP, and AEAp formation was quantified by LC-MS peak area (AUC). ATP supported significantly greater AEAp formation than GTP. Data are mean ± SEM; P value is indicated. **(B)** Divalent cation preference of recombinant human DGKQ. DGKQ was incubated with AEA and ATP in the presence of Mg^2+^, Zn^2+^, Mn^2+^, or the indicated combinations of divalent cations. Mg^2+^ alone supported robust AEAp formation, whereas Zn^2+^ and Mn^2+^ alone supported little or no activity. Data are mean ± SEM; P value is indicated. **(C)** pH dependence of recombinant human DGKQ AEA kinase activity. DGKQ activity was measured across the indicated pH range, with the highest AEAp formation observed at pH 8.5. Data are mean ± SEM.

**Fig. S4.**
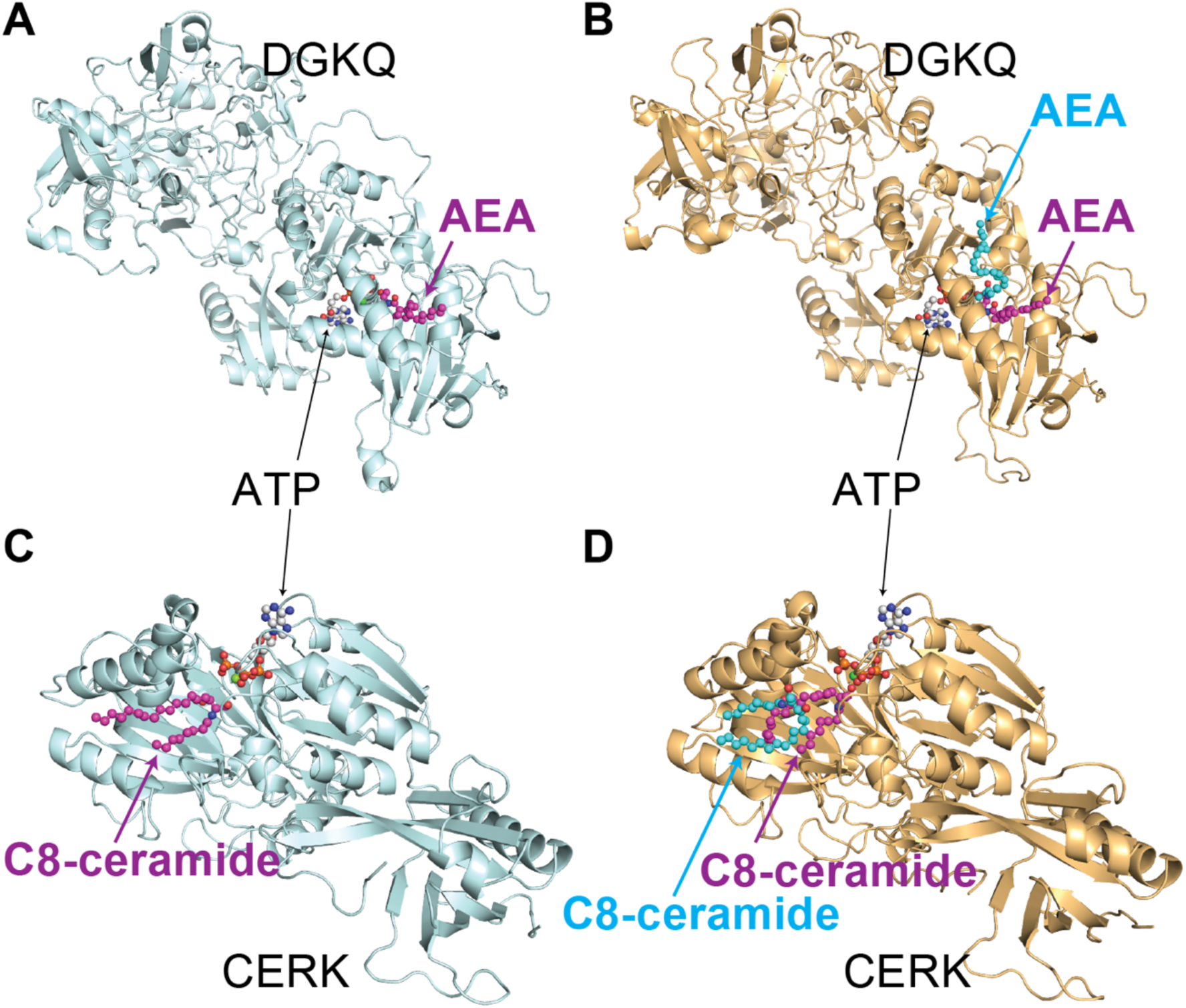
Structural models of DGKQ and CERK bound to one or two lipid substrates. **(A and B)** Boltz-2-predicted structures of DGKQ in complex with ATP and AEA. In the single-substrate model (A), one AEA molecule occupies a hydrophobic cavity adjacent to the ATP-binding site. In the dual-substrate model (B), two AEA molecules can be accommodated simultaneously within an hydrophobic pocket while maintaining proximity to the catalytic center. **(C and D)** Boltz-2-predicted structures of ceramide kinase (CERK) in complex with ATP and C8-ceramide. The single-substrate model (C) shows one C8-ceramide molecule positioned near the catalytic site. In the dual-substrate model (D), two C8-ceramide molecules are predicted to occupy the substrate-binding region, with one positioned near the catalytic site and the other aligned alongside the first molecule. This dual-occupancy model is consistent with previously reported Michaelis-Menten analyses of CERK, in which the substrate-dependence of ceramide phosphorylation appears to deviate from a simple monophasic saturation curve and may be interpreted as biphasic-like behavior (*7*). Thus, the model raises the possibility that simultaneous or sequential occupancy of multiple lipid-binding positions within CERK contributes to non-classical lipid kinase kinetics. The models are presented as structural hypotheses generated by Boltz-2 and have not been experimentally validated.

**Table S1.** EC_50_ values [nM] of canonical LPA ligands and phosphorylated endocannabinoid-related lipids at human LPA receptors.

|  | LPA<sub>1</sub> | LPA<sub>2</sub> | LPA<sub>3</sub> | LPA<sub>4</sub> | LPA<sub>5</sub> | LPA<sub>6</sub> |
| --- | --- | --- | --- | --- | --- | --- |
| **1-oleoyl-LPA** | 19.5 ± 10.6 | 434 ± 262 | 65.9 ± 45.3 | 120 ± 27.8 | 72.2 ± 91.3 | >10 µM |
| **1-linoleoyl-LPA** | 22.5 ± 16.5 | 324 ± 185 | 44.9 ± 31.3 | 111 ± 28.5 | 29.5 ± 28.6 | 1162 ± 372 |
| **2-arachidonoyl-LPA** | 20.1 ± 10.7 | 159 ± 293 | 21.7 ± 41.4 | 15.8 ± 6.47 | 24.7 ± 19.5 | 130 ± 30.5 |
| **OEAp** | 18.0 ± 11.2 | ~4 µM | 622 ± 105 | 192 ± 95.6 | 33.1 ± 22.4 | >10 µM |
| **LEAp** | 43.9 ± 37.4 | ~10 µM | ~7 µM | 455 ± 269 | 27.6 ± 16.5 | ~8 µM |
| **AEAp** | 38.8 ± 29.4 | ~10 µM | ~7 µM | 189.7 | 75.0 ± 128 | ~5 µM |

## Data S1. (separate file)

This file contains the tabulated data behind the figures.

